# Shortlisting genes important in seed maturation by principal component analysis of gene expression data

**DOI:** 10.1101/2020.06.19.158832

**Authors:** Ashish K Pathak, Ridhima Singla, Mamta Juneja, Rakesh Tuli

## Abstract

Transcriptome data are widely used for functional analysis of genes. De-novo assembly of transcriptome gives a large number of unigenes. A large proportion of them remain unannotated. Efficient computational methods are required for identifying genes and modeling those for regulatory and functional roles. Principal component analysis (PCA) was used in a novel approach to shortlist genes, independently of annotation in genome expression data, taking seed development in *Arabidopsis thaliana* as a representative case. PCA was applied to published genome expression data from four lines of Arabidopsis, mutated in seed development. The PC separating all the developmental stages between a mutant and its respective wild type was selected for shortlisting genes as functionally more important. The shortlisted genes identified by PCA belong to a number of biological functions. The genes reported to give sensitivity to desiccation were identified in PCA analysis also in desiccation intolerant lines only. With respect to the network of 98 genes targeted by ABI3, a higher number of genes was identified as important in the mutants abi 3-5, fus 3-3 andlec 1-1 in comparison to abi 3-1. Ontological analysis and comparison with earlier studies suggest that PCA of genome expression data is useful for shortlisting functionally important genes.

## 1. Introduction

Transcriptome studies require shortlisting potentially important genes from a larger set identified in differential expression analysis. Machine learning methods can be useful in such analysis. PCA is an unsupervised machine learning method used to increase the interpretability of gene expression data without loss of information [1]–[3]. It is especially suitable for time series data and allows filtering artifactual variations. Either genes or samples can be taken as the variables[4]. PCA translates the dataset by linear function to find new variables by maximizing the variance, while preserving ‘variability’ of dataset i.e. identifying new variables that can serve as effective predictors. Given a matrix (X) of m variables and n observations, it reduces m variables into r new variables, where r < m. These new r variables account for as much as possible variance explained by m variables while remaining mutually uncorrelated and orthogonal [5], [6].

Transcriptome of three ovule developmental stages in four Arabidopsis lines mutated in transcription factors ABI3 (*abi 3-5*&*abi 3-1*), LEC1 (*lec 1-1*), FUS3 (*fus 3-3*) or LEC2 (*lec 2-1*) were generated and analyzed in an earlier study [7]. These genes are master regulators of seed maturation and desiccation tolerance. Among these, *abi 3-1*&*lec 2-1* resemble in phenotype with the other mutants but are desiccation tolerant. Important genes related to desiccation tolerance were identified, using differential expression analysis. We applied PCA on the publically available [7] gene expression data for identifying stage and mutant-specific important genes. Developmental stages were taken as variables and plotted with different combinations of PCs. We shortlisted important genes using PC that separated the mutant and wild types at all developmental stages. Genes with high differences along the selected PC at each developmental stage of mutant and wild type were considered as more important genes. To check the credibility of the shortlisted genes, the ontologies of the selected genes were compared with the seed maturation process.

During seed formation, seed undergoes a quiescent state. At heart stage; the seed starts preparing itself for the quiescent stage that defines seed maturation. During seed development (early maturation), when embryo grows and seed filling takes place, many seed storage compounds accumulate [8]. At the final stage of seed development (late maturation), the seed acquires desiccation tolerance and becomes quiescent. The transcription factors LEC1, ABI3, LEC2, and FUS3 are the master regulators of seed maturation[9], [10]. Published experimental evidences, like anatomical defects [9, 10], mutant analysis [11], pathway regulation etc. were used in the present study to validate the application of PCA as a gene reduction technique. The analysis suggested that dissimilar sets of gene may be involved in desiccation tolerance modulated by different transcription factors. We examined the relationship among the shortlisted genes to examine if PCA could be applied to gene expression data for shortlisting of functionally more important genes.

## Results

### PCA for shortlisting important genes

Mapping of reads to genome has provided gene expression data for 32,833 genes (including long non-coding RNA) (Additional Data 1). Our analysis shows that the first two components explain 68.2, 65.1, 63.6 and 68.7 percent variance respectively in the data on transcriptome expression in *abi 3-5, abi 3-1, fus3-3* and *lec 1-1* mutants (Table. 1).

**Table 1.**
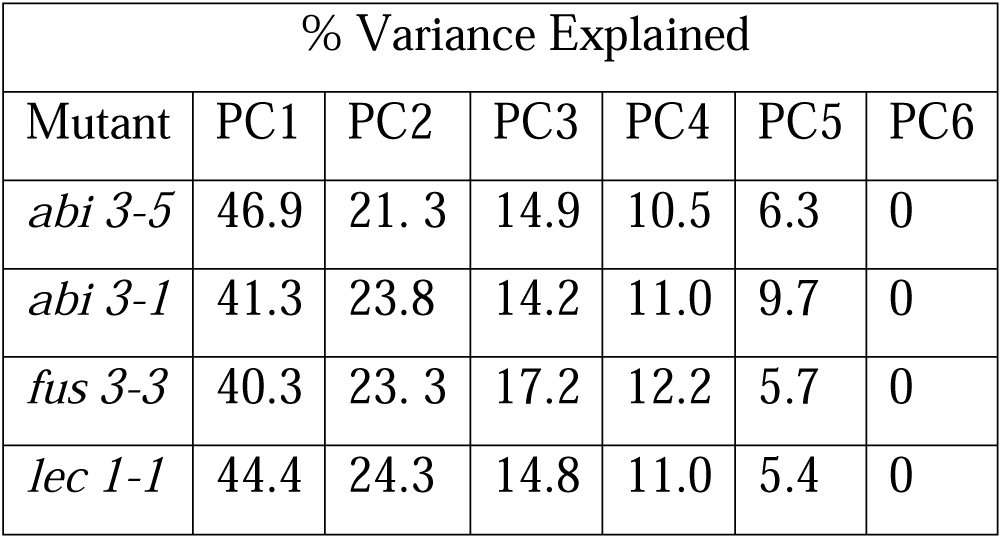
Percentage of variance explained by different principal components in different mutants

| % Variance Explained |  |  |  |  |  |  |
| --- | --- | --- | --- | --- | --- | --- |
| Mutant | PC1 | PC2 | PC3 | PC4 | PC5 | PC6 |
| *abi 3-5* | 46.9 | 21.3 | 14.9 | 10.5 | 6.3 | 0 |
| *abi 3-1* | 41.3 | 23.8 | 14.2 | 11.0 | 9.7 | 0 |
| *fus 3-3* | 40.3 | 23.3 | 17.2 | 12.2 | 5.7 | 0 |
| *lec 1-1* | 44.4 | 24.3 | 14.8 | 11.0 | 5.4 | 0 |

Projections of the six-time points (samples) - three each, for the wild-type and the mutant, were done with different combinations of the PCs. PC1 differentiates genome wide expression in the wild type and the mutants *abi 3-5, fus 3-3* & *lec 1-1* at all three developmental stages (Figure 1). Shortlisting of the relatively important genes for *abi 3-5, fus 3-3* & *lec 1-1* was done along PC1. In case of *abi 3-1*, PC 5 differentiates between the wild type and the mutant (Figure 1). Hence, the shortlisting of important genes for *abi 3-1* was done along the PC 5.

**Figure 1.**
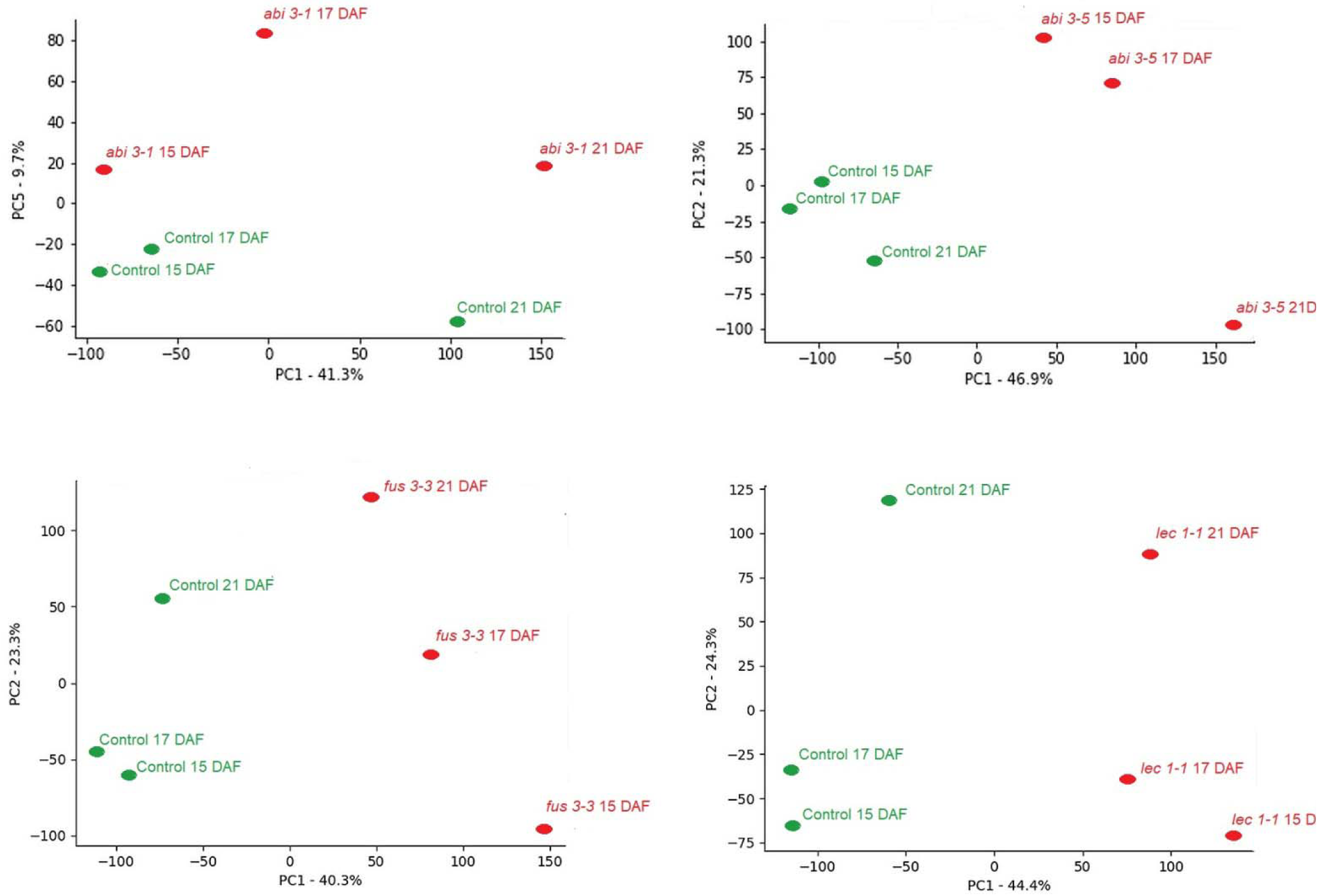
Projection of genomic expression data at different developmental stages along different principal components in *abi 3-5, fus 3-3, lec 1-1* and *abi 3-1* mutants

Genes with a high difference along the identified PC, for each developmental stage between mutant and wild type were identified as the important genes (Additional Data 2). Pair wise comparison of developing ovules revealed common and exclusive genes at 15, 17 and 21 DAF (Figure 2). The highest number of transcripts was noticed in *lec 1-1* at 15 DAF. The *lec 1-1* had the highest number of exclusive genes at 15 DAF, while *abi*3-5 had highest number at 17 DAF, and 21 DAF (Figure 2). Among the important genes identified in different mutants, 956, 500 and 713 were common at 15, 17 and 21 DAF respectively among *abi 3-5, fus 3-3, lec 1-1* and *abi 3-1* (Figure 2). The *abi 3-1* has the lowest number of exclusive genes at all the three developmental stages.

**Figure 2.**
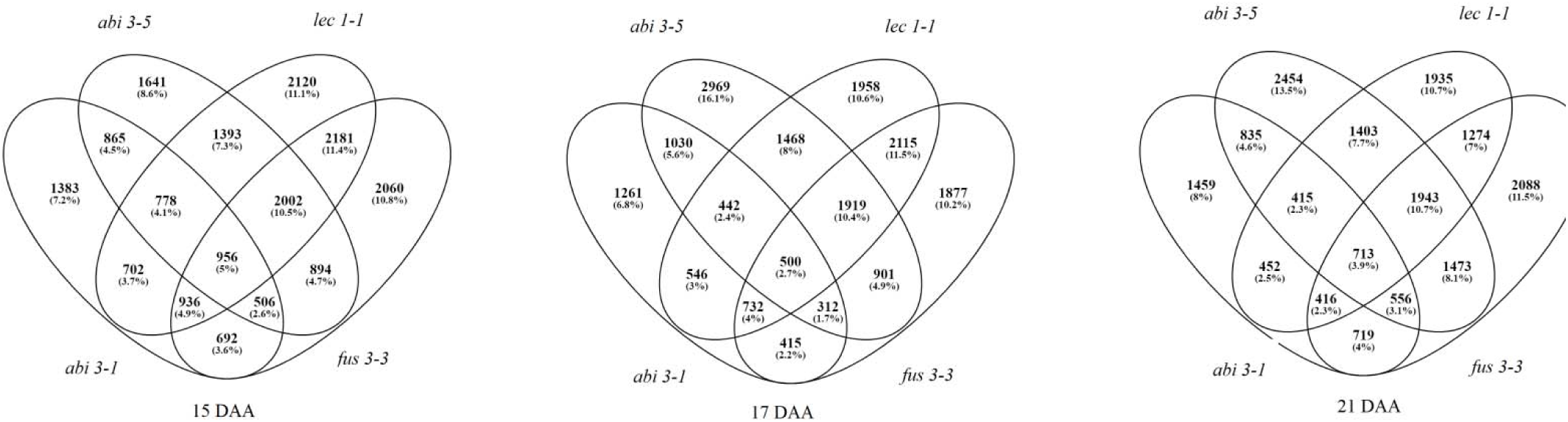
Venn diagram representing the expression of shared and exclusive genes (as unions and intersections) expressed at 15, 17 and 21 DAF in Arabidopsis, to identify functional association of genes in the four mutants.

### Ontologies of shortlisted genes identified in different mutants of seed maturation

During seed maturation, embryo development is arrested and seed dehydrates. The macromolecules that can protect seed viability in dry state are synthesized and stored. LEC1 & FUS3 arrest embryo & endosperm cell division & growth as seed enters dormancy[12]. In the mutant seeds of these genes (LEC1 & FUS3) embryo growth is not arrested&theembryo continues to grow at maturation phase too, and that leads to premature germination of embryo. While ABI 3 does not affect embryo & endosperm growth, it is involved in acquiring dormancy via abscisic acid mediated pathways[13]. The seeds in *abi 3-1* are desiccation tolerant, while those in *abi 3-5, lec 1-1* and *fus 3-3* are dessication intolerant. Reproduction, response to stimulus, metabolic processes, oxidoreductase activity, structural molecule activity and embryonic development ending in dormancy related genes were not shortlisted for *abi 3-1*. Gametophyte development, auxin transport, regulation of multicellular organism development, and response to auxin stimulus were shortlisted only in *fus 3-3*. Trichoblast& root hair cell differentiation related transcripts were shortlisted in *fus3-3*(Additional data 3). The shortlisted genes in *lec 1-1 & fus 3-3*carried the genes for ATP synthesis coupled proton transport, photorespiration, oxidative phosphorylation, photosynthesis & electron transport chain. This leads to the lethality of embryo in these mutants (*lec 1-1*&*fus 3-3*) (Additional data 3). Photosynthesis, thylakoid membrane organization & chlorophyll biosynthetic related genes were shortlisted in *abi 3-5*(Additional data 3).

### Comparison with previous works

We compared the published results on differentially expressed genes associated with seed development in Arabidopsis, with the genes shortlisted in our study. In an earlier report, 2,712 genes were identified as differentially expressed between the desiccation tolerant and intolerant lines. These were classified as desiccation related genes [7]. To examine the reliability of PCA shortlisting method, we compared the distribution of these 2,712 genes in the PCA based shortlisted genes for *abi 3-5, lec 1-1* &*fus 3-3* in comparison with the shortlisted genes for *abi 3-1*. In the three desiccation intolerant mutants i.e. *abi 3-5, lec 1-1* and *fus 3-3*, the distribution of these differentially expressed genes was 6 times higher than that in *abi 3-1*. We identified 908 genes as common between the earlier work and the desiccation related genes identified by PCA (not shortlisted in *abi 3-1*& shortlisted in other desiccation intolerant mutants). OBAP1A, oil bodyassociated protein 1A (AT1G05510), stachyose synthase (AT4G01970), avirulence induced gene 2 like protein (AT5G39720), and DREB2D (AT1G75490), identified as desiccation related genes in the earlier study were also confirmed by the PCA based method for shortlisting. The genes, LEC1, FUS3 & ABI3 are known to influence the expression of one another[14]. In our analysis also, ABI3 (AT3G24650) was shortlisted for influencing both FUS3 & LEC1, LEC1 for ABI 3 (in both *abi 3-1*&*abi 3-5*) & FUS3. However, FUS3 was shortlisted by PCA only in LEC1. In an earlier work[15], a set of 98 genes was suggested as the targets regulated by ABI3. These constitute the regulatory network expressed during seed maturation. We compared our results to examine which of the targets of ABI3 were shortlisted in the PCA based analysis also. Combining the three developmental stages, 65, 38, 35 and 24ABI regulated genes were identified respectively in *abi 3-5, fus 3-3, lec 1-1* and *abi 3-1*mutant lines (Figure 3). The *abi 3-5* has the highest percentage of ABI regulated genes across alldevelopmental stages.Among the targets of ABI 3 identified in different mutants (combining all three developmental stages), 17 genes were identified in *abi 3-5, fus 3-3* and *lec 1-1*, while these were absent in *abi 3-1*&had been identified as desiccation related genes in the previous report.Our results also support that these 17 genes may be the candidate genes involved in desiccation tolerance (Additional data 4).

**Figure 3.**
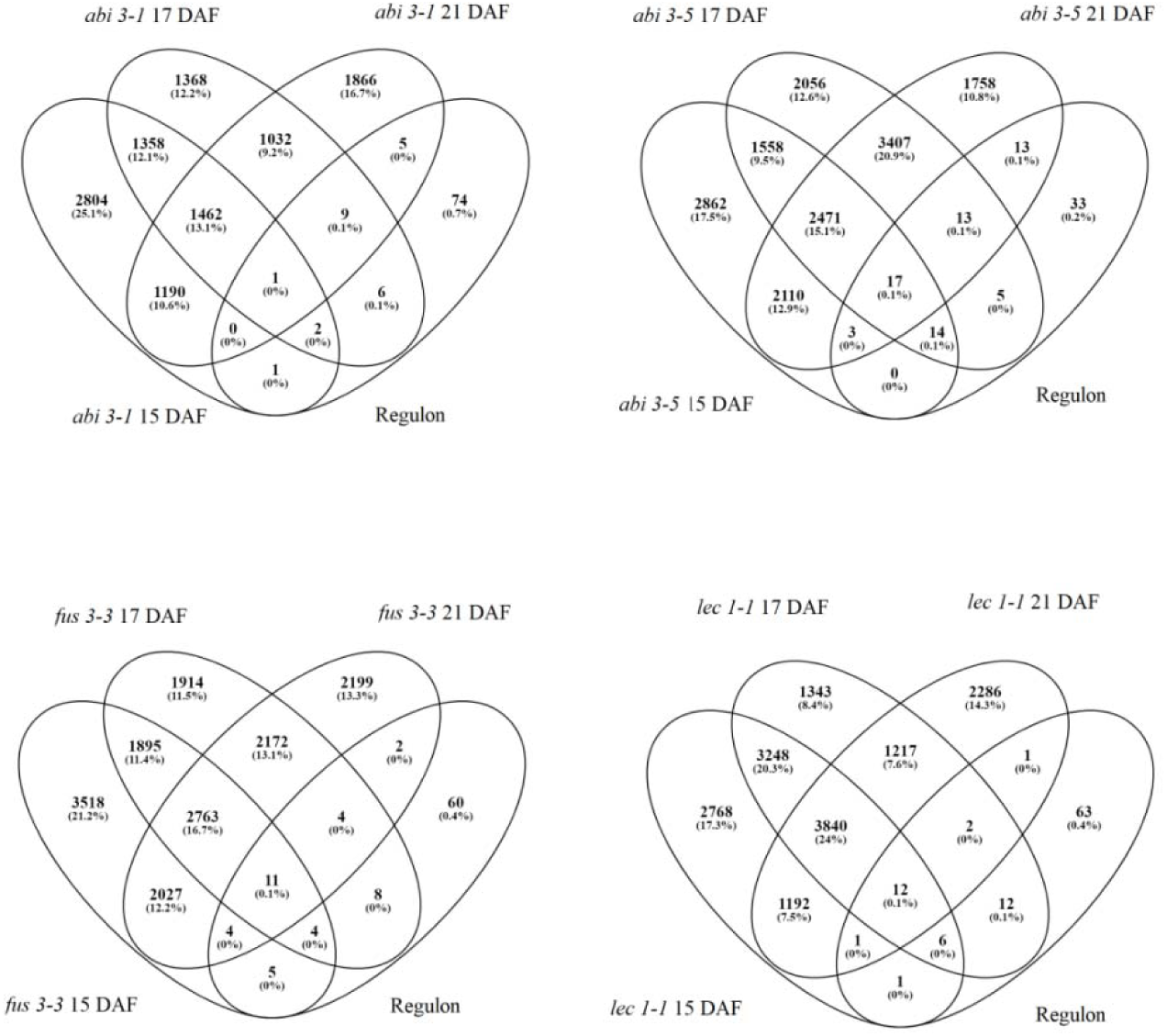
Venn diagram representing the expression of shared and exclusive genes identified as more important by PCA multivariate analysis at three ovule developmental stages in the wild-type, *abi 3-5, fus 3-3, lec 1-1* and *abi 3-1* mutants

### Long non-coding RNA

Long non-coding RNA are transcripts with length more than 200 nucleotides that do not code for proteins. Theseplay important role in regulation of seed development[16]. In *abi 3-1, abi 3-5, fus 3-3* and *lec 1-1*, 854, 868, 1024 & 1023 genes were shortlisted respectively (Additional data 5). Long non coding RNAs involved in RNA splicing were present in desiccation intolerant mutants and absent in *abi 3-1*. This suggests, RNA splicing by long coding RNA might be involved in determining desiccation tolerance. Interestingly, all the genes represented in RNA splicing in *abi 3-5* were completely different from *lec 1-1* and *fus 3-3*. Between *lec 1-1*&*fus 3-3*, except two unique genes, others were the same. Independent ontology analysis of long coding RNA was also in agreement, suggesting that overlapping but not redundant pathways control seed dormancy by LEC1, FUS3 & ABI3 (Additional data 5; Table 2).

**Table 2.**
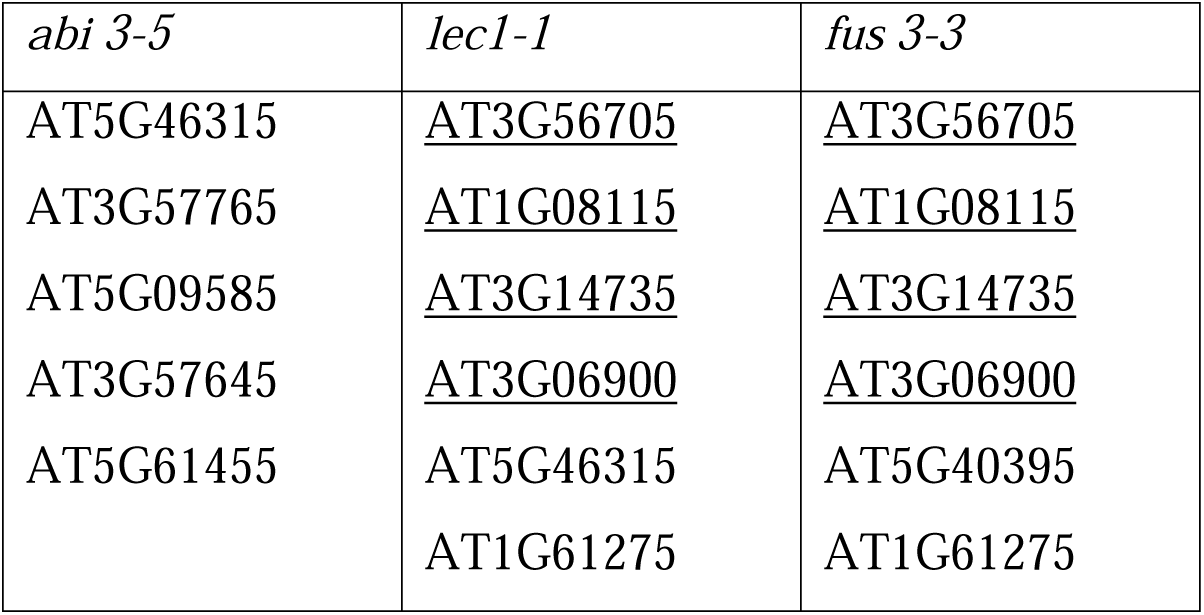
RNA splicing related long non coding RNA genes shortlisted in *abi 3-5, lec1-1* and *fus 3-3*.

## Discussion

Shortlisting of functionally more important genes is the first step in the interpretation of high throughput gene expression data. Normalization of gene expression data is a commonly used technique to make samples comparable. However, the normalization steps do not correct the artifacts such as batch effect in gene expression analysis. Errors in the selection of trait-related genes can be minimized by multiple approaches [4], [17], [18]. One of the promising methods to resolve these artifacts is to examine the PCs of the data.

PCA allows the visualization of multidimensional data by representing it along new dimensions (PC). It is also used as a clustering method (Figure 1). The first few PCs may not necessarily capture most of the cluster structures [5]. PC5 was able to separate all the developmental stages between the mutants and wild type (Figure 1) in *abi 3-1*. PCA is performed by considering either genes [3], [19] or the experimental conditions as the variable [20], [21]. Genes asthe variable, create PCs that indicate characteristics of genes which best explain the experimental response they produce. When experimental conditions are the variable, these create a PC that indicates characteristics of experimental conditions as explaining gene behavior they elicit. Various analyses have been reported, considering experimental conditions as the variables, as in case of sporulation time series gene expression data on yeast [4]. Suitability of sample embedded PCA for shortlisting genes is well described for time series gene expression data [21]. PC that separates all developmental stages of the mutant and wild type is interpreted to capture variance explaining phenotypic differences between the mutant and wild type (Figure 1). These variables (developmental stages) in Figure 1 represent the summation of all the gene products, using scaled gene expression with its loading score. Multiplication of the “scaled CPM” with “loading score” provides co-ordinate of gene along the PC. Genes with high difference along the identified PC, between the developmental stages of wild type and mutants were classified as potentially more important genes. Genes identified by this method were compared with published literature to verify correctness of the approach, as applied to seed maturation (Additional data 1 & 2).

Biological results are difficult to capture and convey in mathematical relationships. Ontologies provide a structured representation of knowledge gained by various approaches. Genes shortlisted by PCA were rich in the processes involved in seed maturation. Enrichment of reproduction, response to stimulus, metabolic processes, oxidoreductase activity, structural molecule activity and embryonic development ending in dormancy related genes in *abi 3-5,lec 1-1* and *fus 3-3*, while not in *abi 3-1*agrees with the normal cotyledon in *abi 3-1*. Shortlisting of gametophyte development, auxin transport, regulation of multicellular organism development, and response to auxin stimulus only in *fus 3-3*, agrees with the involvement of FUS3 in cell growth arrest of embryo[7]. Trichoblast & root hair cell differentiation related transcripts were shortlisted in *fus3-3*(Additional data 3),FUS 3 are reported form abnormal embryo root apices[22]. The embryo lethality of *lec 1-1*&*fus 3-3* agrees with shortlisting of ATP synthesis coupled proton transport, photorespiration, oxidative phosphorylation, photosynthesis & electron transport chain in *lec 1-1 &fus 3-3*(Additional data 3). The ontology of shortlisted genes agrees with different pathways that control seed maturation in LEC1 & FUS3 [23]. Shortlisting of photosynthesis, thylakoid membrane organization & chlorophyll biosynthetic related genes were shortlisted in *abi 3-5*, in agreement with chlorophyll retention in *abi 3-5*[11](Additional data 3).

Earlier report [7] suggested that the genes down-regulated in desiccation intolerant mutants (*lec 1-1, fus 3-3* and *abi 3-5*) as compared to desiccation tolerant lines (*abi 3-1* and *lec 2-1*) are the desiccation related genes. The distribution of dessication related genes identified by differential expression analysis[7].establishes the reliability of our approach. Shortlisting of OBAP1A, oil bodyassociated protein 1A (AT1G05510), stachyose synthase (AT4G01970), avirulence induced gene 2 like protein (AT5G39720), and DREB2D (AT1G75490) as dessication related genes by both differential expression analysis and PCA based shortlisting suggest high probability of their involvement in acquiring dessication tolerance in seeds. The presence of exclusive sets of PCA shortlisted genes among studied transcription factors agrees with the involvement of different sets of genes regulated by those transcription factors[12], [13]. Genes shortlisted by PCA agree with cross regulation among the studied transcription factors [24], [25]. Genome-wide chromatin immunoprecipitation (ChIP-chip), transcriptome analysis, quantitative reverse transcriptase– polymerase chain reaction and transient promoter activation assay were combined to identify a set of 98 ABI3 target genes[15]. We compared the genes shortlisted by PCA with the ABI 3 targeted genes. A large number of ABI 3 targets was present among the genes shortlisted by PCA in *lec 1-1, fus 3-3* and *abi 3-5* (Figure 3). In *abi 3-1* least number of such targets was identified. This agrees with the leaky phenotype of *abi 3-1*. Targets of ABI 3 not shortlisted by PCA in *abi 3-1*, were present in the other three mutants. These may be important targets for desiccation tolerance. The presence of the highest number of ABI 3 targets in *abi 3-5*further supports the accuracy of PCA based shortlisting of genes. In this study, we used gene expression data at 15, 17 and 21 DAF, while [12] performed transcriptome analysis at 5, 7, 9, 11, 13, 15, 17 and 19 days after flowering. The unidentified ABI 3 targets may be involved at early stages.The absence of the indicators of RNA splicing in *abi 3-1*, and their presence in other three mutants, suggestthatdessication tolerance might be regulated by RNA splicing. ABI 3 splicing is known to determine seed dormancy[26].

## Conclusions

PCA has previously been used for shortlisting genes by selecting high and low scoring genes along different PCs. Our method differs from others, as we have shortlisted functionally important genes on the basis of PC that separates wild type and mutants at multiple stages of seed development. Co-ordinates of the genes were calculated at each developmental stage for wild type and mutants along the identified PC. Genes with a higher differencebetween the mutant and wild typeat different developmental stages (15 DAF, 17 DAF& 21 DAF), along the selected PC were shortlisted. Our study provides an easily understandable application of PCA to analyze time series gene expression data. Exploring time series relationships can provide good basis to simulate the dynamic gene regulatory network. Clustering methods such as DTW and the variants of PCA need to be further examined for resolution of the time series patterns.

## Methods

The data was downloaded from EBI-ENA (https://www.ebi.ac.uk/ena) in fastq format[7]. The genome of *Arabidopsis thaliana* was obtained from ensembl database (https://plants.ensembl.org/index.html)and indexed using Bowtie build at default parameters. The raw reads were mapped using Bowtie version 2.1.0-2[27] and stored as bam format. The bam file were sorted and indexed using samtools[28]. Read counts were obtained using HTSeq[29] and normalized as counts per million (CPM) using edgeR[30].

Gene expression data matrix with 6 variables (columns) i.e. three developmental stages (15, 17 and 21 DAF) from the mutant and wild type wasstored in CSV format. The CSV file was loaded and stored in python using pandas module. To compute principal PCs, eigenvalues and loading score, and for plotting the graphs, available algorithms in different libraries like Sklearn and Matplotlib were used.

Different developmental stages were considered as variables and the genes were taken as observations. The variables (developmental stages) were plotted along different combinations of PCs in a set of two. The PC separating themutant and the wild type at all developmental stages was used for shortlisting the genes. On the basis of the results of PCA, the observations i.e. genes were reduced by identifying genes with a high difference (day wise comparison between mutant and wild type) along with PC explaining the variance between the wild type and mutants at all the developmental stages. The loading score for each gene along with the identified PC for various combinations of mutants was calculated. The scaled CPM values for all the genes were calculated for four different mutants. These values were multiplied with their loading score to CPM after scaling. The absolute difference in the values at each developmental stage between the mutant and wild type was calculated. The average of the absolute difference of all the genes was taken as the threshold, and the genes having values above the threshold were selected as the important genes.

Statistical test for enrichment of ontologies to check the trustworthiness of the shortlisted genes by PCA was used from AgriGO web-based tool (http://bioinfo.cau.edu.cn/agriGO/) using singular enrichment analysis (SEA) method.

## Additional files

**Additional Data 1**. Gene expression data for *abi3-1, abi 3-5, fus 3-3*&*lec 1-1* with their respective control at three developmental stages of seed.

**Additional Data 2**. Functionally more important genes shortlisted by PCA, for *abi 3 - 1, abi3-5, fus3-3*&*lec1-1*& at 15, 17 and 21 DAF.

**Additional Data 3**. Enriched ontologies identified in mutants at different developmental stages on the basis of molecular function (sheet 1) & enriched ontologies identified in mutants at different developmental stages on the basis of biological function (sheet 2).

**Additional Data 4**. List of candidate genes identified for desiccation tolerance on the basis of differential expression analysis from previous work & PCA based shortlisting method

**Additional Data 5**. Functionally more important long non-coding RNA shortlisted by PCA, for *abi3-5, fus3-3, lec1-1*&*abi 3-1* at 15, 17 and 21 DAF.

## Abbreviations

DAF: Days after flowering
PCA: Principal component analysis
PCs: Principal Components
ABI3: Abscisic acid insensitive 3
LEC1: Leafy cotyledon 1

## Acknowledgements

RT and AKP acknowledge Science & Engineering Council of Department of Science & Technology, Govt of India for JC Bose Fellowship to RT and MHRD for support under Design Innovation Centre, Govt of India.

## Availability of data and materials

The Python implementations are available at https://github.com/ridhima-singla/PCA_Arabidopsis-

## Authors’ contributions

R.T. proposed and designed the study. A.K.P and R.S performed the experiments, and analyzed data. M.J guided the experiments and analysis. A.K.P. wrote the manuscript, R.T. improved presentation. All authors read and approved the manuscript.

## Ethics approval and consent to participate

Not applicable.

## Consent for publication

Not applicable.

